# NOD2 stimulation reduces *Cryptosporidium parvum* level of infection in neonates by promoting intestinal epithelial production of antimicrobial peptides and cell renewing

**DOI:** 10.1101/2025.03.14.643285

**Authors:** Mégane Fernandez, Tiffany Pezier, Julien Pichon, Yves Le Vern, Catherine Werts, Sonia Lacroix-Lamandé

**Author notes:** Corresponding author. (SLL).

## Abstract

At birth, the mucosal immune system of neonates is not fully developed, making them more susceptible to respiratory and intestinal diseases. Previously described host-directed therapies using TLR activation-based strategies have proven effective in controlling neonatal diseases, including cryptosporidiosis. In this study, we investigated the effect of NOD receptor stimulation on the control of enteric infection by the protozoan *Cryptosporidium parvum* in neonatal mice. NOD2 stimulation by intraperitoneal injection of MDP resulted in a rapid reduction in the parasite burden. The protective effect was associated with increased proinflammatory cytokine and antimicrobial peptide gene expression and a rapid influx of neutrophils to the site of infection, whereas NOD1 stimulation did not show a protective effect. The protective mechanism did not involve microbiota participation but involved IFN-γ and IL-22 cytokines and was associated with increased intestinal epithelium renewal in infected neonates. Our findings, demonstrated in neonatal mice, elucidate how the stimulation of NOD2 with the bacterial ligand MDP actively contributes to the non-specific clearance of a parasitic infection within the intestinal epithelial cells.

**Author Summary:** Newborns are particularly vulnerable to intestinal infections due to their immature immune systems. Developing strategies to enhance their ability to fight infections is crucial for improving neonatal health. In this study, we investigated how stimulation of a specific immune receptor, NOD2, can help neonatal mice control an intestinal infection caused by the protozoan Cryptosporidium parvum. We found that intraperitoneally administration of a bacterial molecule called MDP, which activates the NOD2 receptor, led to a rapid reduction in the parasite burden. This protection was associated with increased production of inflammatory molecules, antimicrobial peptides, and a significant influx of neutrophils to the site of infection. Importantly, the protective effect did not depend on the gut microbiota but involved specific immune signals, such as IFN-γ and IL-22, and enhanced the renewal of intestinal epithelial cells. These findings highlight how targeting NOD2 can boost neonatal innate immunity, offering a potential host-directed strategy to combat severe intestinal infections, limit long-term intestinal damage and reduce the risk of fatal outcomes.

## Introduction

Cryptosporidiosis is an intestinal disease caused by the protozoan *Cryptosporidium parvum* (*C. parvum*), typically transmitted through oral ingestion. This parasite develops in intestinal epithelial cells (IECs) and causes acute diarrhea in very young ruminants, young children and immunocompromised hosts (1, 2). The hallmark of this disease is acute gastrointestinal distress manifested by diarrhea, abdominal cramps, nausea, vomiting, and fever. In humans, the most susceptible groups include children under the age of 5 and immunocompromised individuals (2). Importantly, *C. parvum* is the second most common cause of moderate to severe diarrhea among children under the age of 2 (3). In contrast, in cattle, cryptosporidiosis is recognized as endemic worldwide and is one of the most important causes of neonatal enteritis in calves, resulting in substantial financial losses for livestock farmers (1).

Susceptibility to this infection is intricately linked to the host immune response, highlighting the central role of the immune system in combating this parasitic invasion. We and others and have highlighted the critical role of innate immunity in controlling the acute phase of *C. parvum* infection, whereas adaptive immunity is required for definitive clearance of the infection (4, 5). Host epithelial cells play a major role in both the multiplication of the parasite and the initiation of immune responses by recruiting inflammatory immune cells such as neutrophils and dendritic cells to the site of infection (4, 6–8). In addition, conventional type-1 dendritic cells (cDC1) play a major role in this innate protective response (9) whereas inflammatory monocytes rapidly recruited to the site of infection promote the loss of intestinal barrier integrity that characterizes cryptosporidiosis (10).

With no vaccine available and limited chemotherapy, there is an urgent need to develop new methods to control cryptosporidiosis. Considering the close relationship between the susceptibility of this infection and the immune status of the host, the exploration of immune-enhancing strategies is a promising avenue to consider. Our hypothesis is that the enhancing innate immunity with potent immunostimulants, such as pattern recognition receptor (PRR) ligands, could stimulate the production of key cytokines, that may provide defense against this enteric pathogen. In the neonatal mouse model, we have previously shown that stimulation of innate immunity by Toll-like receptor (TLR) agonists, including CpG-ODN (TLR9 ligand), or PolyI:C (TLR3 ligand), increased the intestinal immune responses and subsequently reduce the intestinal burden of the *C. parvum* parasites (11, 12).

Another class of PRRs known as nucleotide-binding oligomerization domain (NOD) receptors plays a critical role in the function of the innate immune system (13–16). NOD1 and NOD2 belong to this intracellular PRR family and recognize different fragments of peptidoglycan (PGN), a major component of the bacterial cell wall (17). NOD1 recognizes γ-D-glutamyl-mesodiaminopimelic acid (iE-DAP) from Gram-negative bacteria and certain Gram-positive bacteria and NOD2 recognizes muramyl dipeptide (MDP), which is present in both Gram-positive and Gram-negative bacteria. Several genome-wide association studies have identified NOD2 as the most prevalent susceptibility factor for the development of Crohn’s disease, highlighting the importance of NOD2 in intestinal immunity (16, 18, 19). In the intestine, NOD2 is expressed by both hematopoietic and non-hematopoietic cells that constitute the intestinal epithelium (20–24). Upon activation by MDP, NOD2 facilitates host defense mechanisms by triggering the production of cytokines, chemokines, antimicrobial peptides, and mucins (20, 25–27). Several studies have described the protective role of NOD2 in various models of intestinal inflammation (17, 27, 28). Furthermore, activation of NOD2 has been shown to improve clearance and reduce the severity of *Salmonella* and *Citrobacter rodentium* colitis in mice, highlighting its involvement in the regulation of the intestinal immune response (29, 30). In addition, NOD2 activation has been shown to ameliorate intestinal bacterial infections caused by *L. monocytogenes*, *S. flexneri*, and *Helicobacter hepaticus*, primarily through the production of pro-inflammatory mediators, antimicrobial peptides and bacterial autophagy (17, 26, 31).

In this study, we investigated the role of NOD stimulation in the regulation of *C. parvum* infection in neonatal mice. Our findings demonstrate that the stimulation of NOD2 by the MDP agonist promotes a protective effect by enhancing the innate immune response, leading to the renewal of the epithelial barrier.

## Results

### Stimulation of NOD2 receptor reduce *C. parvum* intestinal load in infected neonatal mice

We previously reported that stimulation of TLR9 and TLR3 receptors in neonatal mice can reduce the severity of cryptosporidiosis by enhancing a protective innate immune response in the intestine (11, 12). While the signal transduction pathways differ between TLR and NOD receptors upon binding of their respective agonists, we asked whether NOD receptor stimulation could elicit also a protective effect against *C. parvum* infection in neonatal mice by using the *γ*-D-glutamyl-*meso*-diaminopimelic acid (iE-DAP) and the muramyl dipeptide (MDP) as NOD1 and NOD2 agonists, respectively. To investigate this, neonatal mice of 3-day-old were inoculated orally with 5x10^5^ oocysts of *C. parvum* and five days later received 200µg of NOD agonists orally or intraperitoneally (IP) (Fig 1A). Mice were sacrificed the following day to assess the intestinal parasite burden.

**Fig 1.**
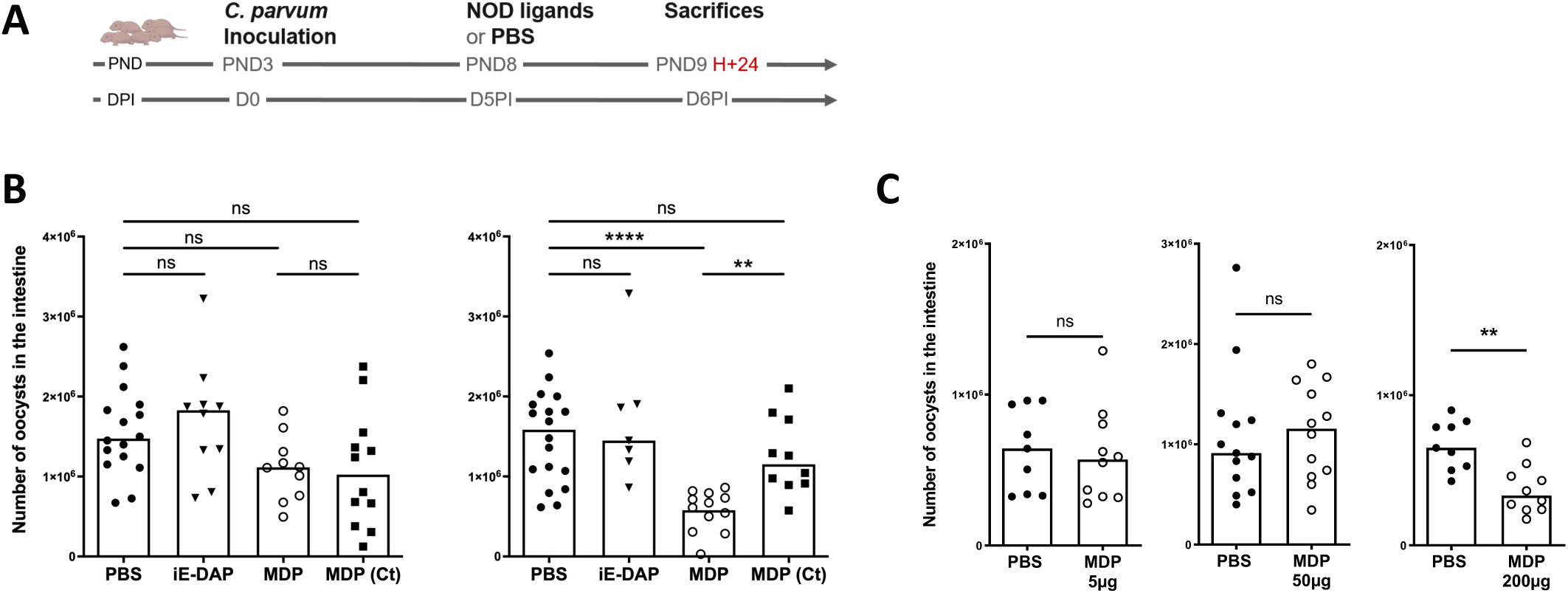
NOD2 stimulation reduces *C. parvum* infection in neonatal mice. (**A**) Experimental timeline. PND3 neonatal mice were orally infected with 5.10^5^ oocysts of *C. parvum* and received 200µg of the NOD-ligands at day 5 p.i.. (**A**). iE-DAP (NOD1 ligand), MDP (NOD2 ligand) and its control MDP (MDP (Ct)), or PBS (control group) were administered by intraperitoneal or oral route to neonatal mice. (**B**) Parasite load in the intestine was evaluated 24 h later (n = 10–18 mice for each group). Statistics are calculated by the one-way Kruskal-Wallis test. (**C**) MDP was administered at 5µg, 50µg and 200µg (n = 9–12 mice for each group). Statistics are calculated by the Mann-Whitney test. Each point corresponds to one mouse, and bars represent the median of each group. ns>0.05, **P<0,01, ****P<0,0001. PND = postnatal days; DPI=days post-inoculation.

As shown in Fig 1B, no reduction in parasite load was observed after oral NOD1 or NOD2 stimulation. In contrast, IP administration of NOD2 agonist (MDP) significantly reduced the parasite load in infected neonatal mice whereas NOD1 stimulation (iE-DAP) was not effective. Notably, a high dose of the NOD2 ligand was required for this protection, as 5 and 50µg of MDP administered under similar conditions did not decrease infection levels (Fig 1C).

### Increased systemic and intestinal inflammatory responses in neonatal mice intraperitoneally injected with MDP

We have previously shown that a transient increase in the inflammatory response induced by TLR stimulation can help to reduce *C. parvum* infection in neonatal mice (11, 12). Here, upon MDP injection by IP route into *C. parvum*-infected neonatal mice, a significant increase in the percentage of neutrophils (CD45+ CD11b+ Ly6G+) was observed in the peritoneal cavity 4 h after injection, whereas the percentage of macrophages (CD45+ CD11c+ MHCII+ CD64+) and DC (CD45+ CD11c+ MHCII+ CD64-) was significantly decreased (Fig 2A). This shift of immune cell percentage was associated with a significant increase in the gene expression of some pro-inflammatory cytokines and chemokines (IL-1β, IL-6, IL-12p40, CXCL1 and CXCL2) in total of intraperitoneal cells (Fig 2B), with the exception of TNF-α expression being decreased. This increased inflammatory response was also observed at the systemic level within the spleen, exhibiting a substantial increase in IL-1β, IL-6, TNF-α, CXCL1 and CXCL2 (Fig 2C). In the ileum tissue, at the site of infection of *C. parvum,* similar to the peritoneal cavity and the spleen, MDP injection significantly increased the gene expression of IL-1β, IL-6, TNF-α, IL12-p40, CXCL1 and CXCL2 (Fig 3A) and also in purified IECs (Fig 3B). Twenty-four hours after MDP injection, these modifications of gene expression were no more visible (Figs S3-A and S3-B).

**Fig 2.**
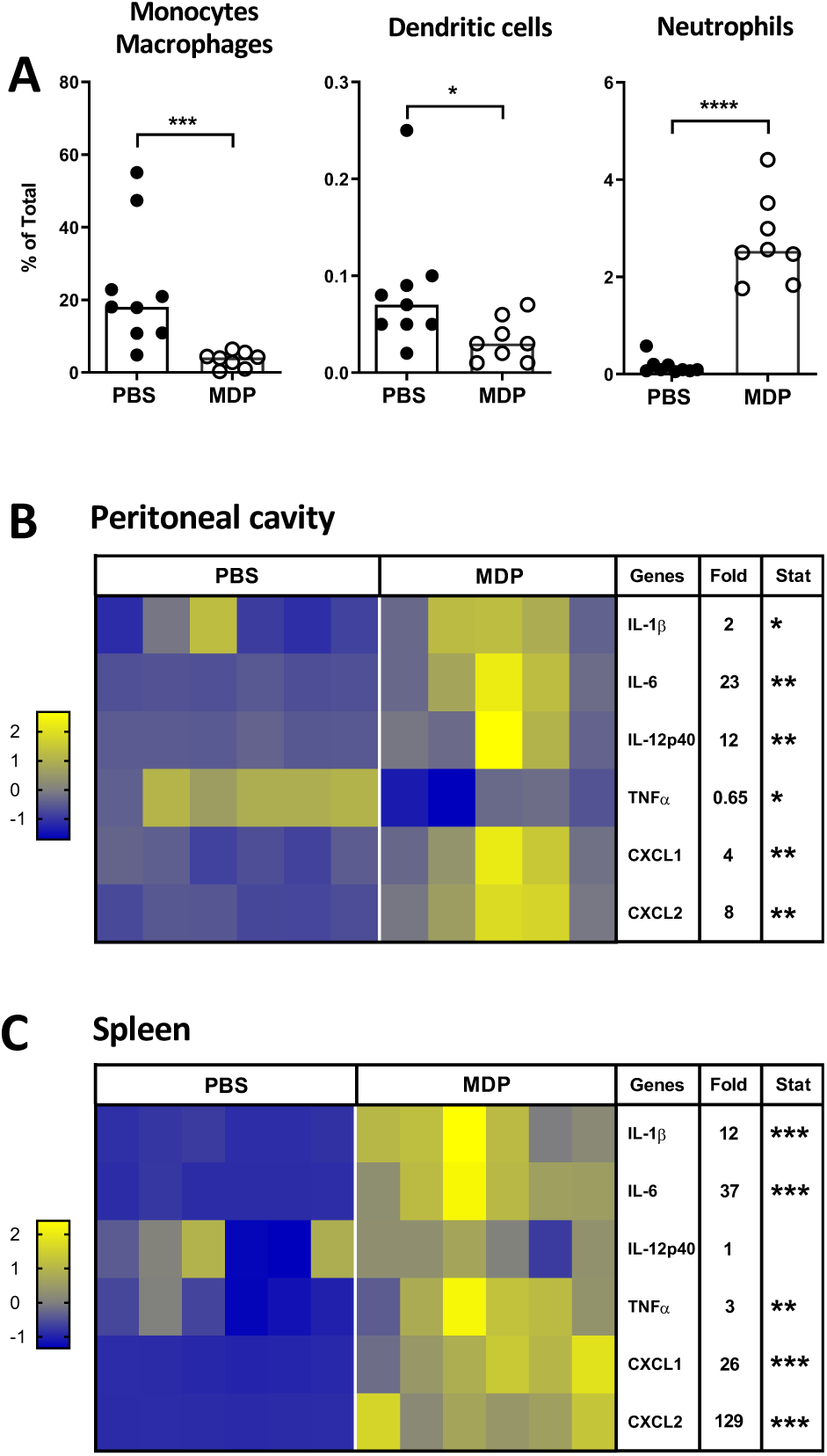
MDP injection into *C. parvum*-infected-neonatal mice increases the inflammatory response in the peritoneal cavity and in the spleen. PND3 neonatal mice were orally infected with 5.10^5^ oocysts of *C. parvum* and received 200µg of MDP by IP route at day 5 p.i. After 4 h, cells from the peritoneal cavity were collected and analyzed by flow cytometry (**A**) and by qRT-PCR (**B**). The percentage of monocytes/macrophages CD45+ CD11c+ MHCII+ CD64+, dendritic cells CD45+ CD11c+ MHCII+ CD64-, neutrophils CD45+ Ly6G+ CD11b+ was determined (n = 8–9 mice for each group). After 4 h, spleens were sampled and analyzed by q-RT-PCR (**C**). (**B-C**) Levels of pro-inflammatory gene expression were quantified by RT-qPCR. Heat maps are designed by z-score of the 2^-Δct^ results for each gene and fold change of mRNA expression calculated as 2^-Δct^ results of the MDP-group in comparison to the PBS-group (n = 5-6 mice for each group). Statistics are calculated by the Mann-Whitney test. ns>0.05, *P<0.05, **P<0,01, ***P<0,001, ****P<0,0001. PND = postnatal days.

**Fig 3.**
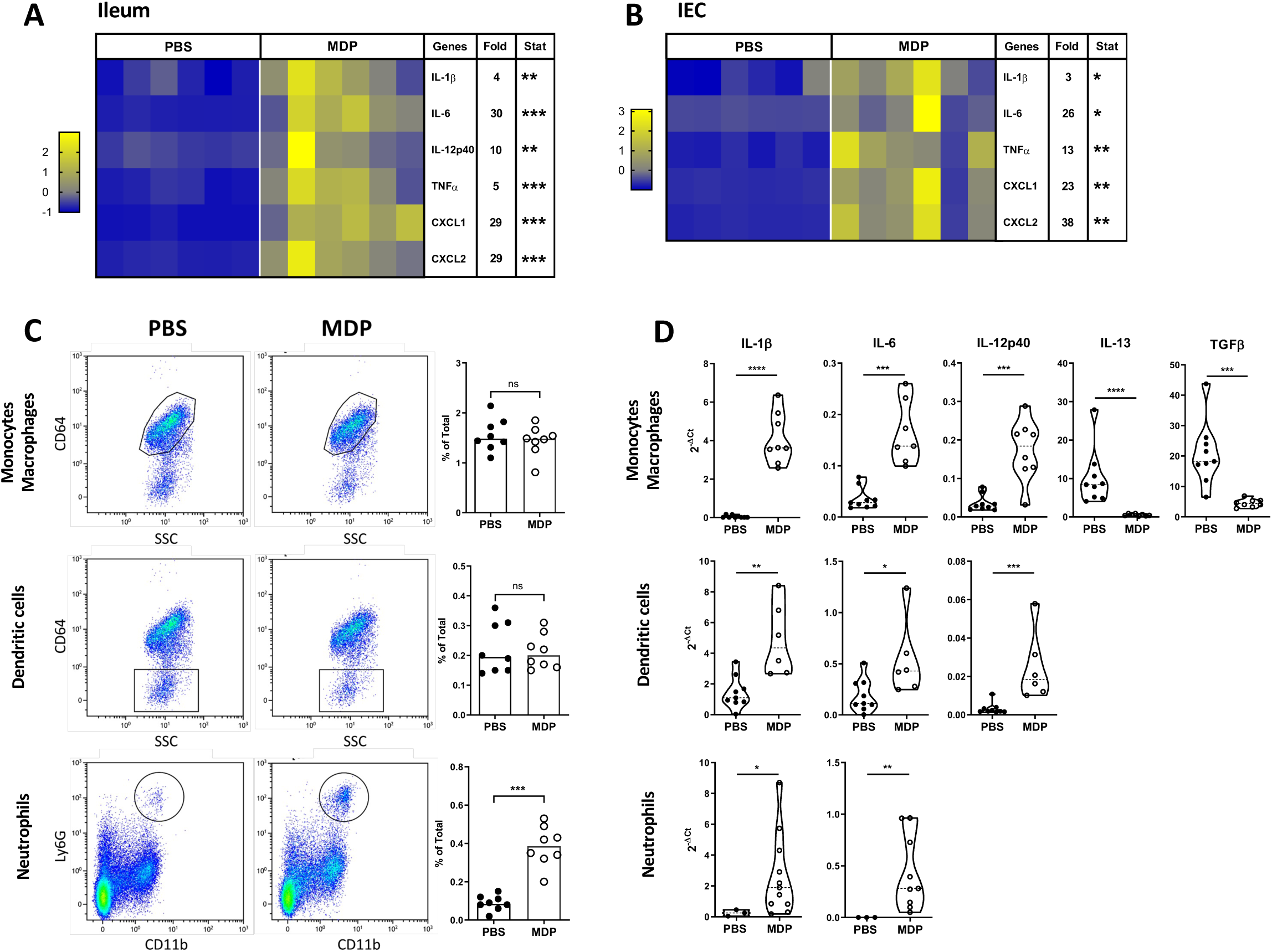
The systemic injection of NOD2 agonist increases inflammation in the ileum of *C. parvum*-infected-neonatal mice. PND3 neonatal mice were orally infected with 5.10^5^ oocysts of *C. parvum* and received 200µg of MDP by IP route at day 5 p.i.. Four hours after MDP injection, the intestine was sampled. The levels of inflammatory gene expression were quantified by RT-qPCR in the ileum tissue (**A**) and in purified IECs (**B**). Heat maps are designed by z-score of the 2^-Δct^ results for each gene and fold change of mRNA expression calculated as 2^-Δct^ results of the MDP-group in comparison to the PBS-group (n = 6 mice for each group). (**C**) Cells from the *lamina propria* of the distal small intestine were collected and analyzed by flow cytometry. The percentage of monocytes/macrophages CD45+ CD11c+ MHCII+ CD64+, dendritic cells CD45+ CD11c+ MHCII+ CD64-, neutrophils CD45+ Ly6G+ CD11b+ was determined. (n = 8 mice for each group). (**D**) Monocytes/macrophages, dendritic cells and neutrophils were sorted, and the levels of inflammatory gene expression were quantified by RT-qPCR. Results are expressed as the 2^-Δct^ results for each gene and compared between the MDP-group and the PBS-group (n = 9-11 mice for each group). Statistics are calculated by the Mann-Whitney test. ns>0.05, *P<0.05, **P<0,01, ***P<0,001, **** P<0,0001. PND = postnatal days.

The inflammatory state of the intestine after MDP injection was evident by neutrophils recruitment as early as 4 h and was still visible 24 h after injection (Figs 3C and S3-C). In contrast, no changes were observed in the percentage of monocytes/macrophages (CD45+ CD11c+ MHCII+ CD64+) and of dendritic cells (CD45+ CD11c+ MHCII+ CD64-) (Fig 3C). Sorting neutrophils, monocytes/macrophages, and dendritic cells from the ileum of infected neonatal mice revealed increased mRNA gene expression of IL-1β and IL-6 in each cell population after MDP injection (Fig 3D). Increased IL-12p40 mRNA gene expression was observed in dendritic cells and monocytes/macrophages from MDP-injected neonatal mice. In addition, the monocytes/macrophages expressed less IL-13 and TGFβ mRNAs (Fig 3D). These results may indicate a shift in resident macrophages toward a pro-inflammatory profile, characteristic of “M1” macrophages. Altogether, these data demonstrate that MDP injection into neonatal mice stimulates a rapid and transient inflammatory response both at the injection and infection sites as well as in the spleen.

### Impact of MDP injection on intestinal epithelial cells in neonatal mice

NOD2 receptor is expressed by various immune cells and IECs (20) and its stimulation is known to trigger the secretion of the antimicrobial peptides. Significant overexpression of the antimicrobial peptides Reg3β/γ and S100A8/A9 genes was observed in the IECs of MDP-injected neonatal mice compared to PBS-injected neonatal mice (Fig 4A). These data suggest that IECs may directly respond to MDP, as described previously with intestinal organoid models (32), or to the inflammatory environment induced by MDP, thus contributing to protection. Considering the modification in antimicrobial peptide responses after MDP injection, one may hypothesize that microbiota could play a role in reducing the intestinal parasite load as previously described after PolyI:C injection (12). To investigate this, we used an experimental model in which pregnant mice received a mixture of broad-spectrum antibiotics in their drinking water 2 days before the expected day of delivery and throughout the experiment in order to reduce the bacterial loads in their intestines and that of their pups (Fig 4B). However, upon administration of MDP to neonates born from antibiotic-treated dams, the reduction in the parasite load was not significantly modified compared to conventional neonates, suggesting that the microbiota is not involved in the protection induced by MDP (Fig 4C).

**Fig 4.**
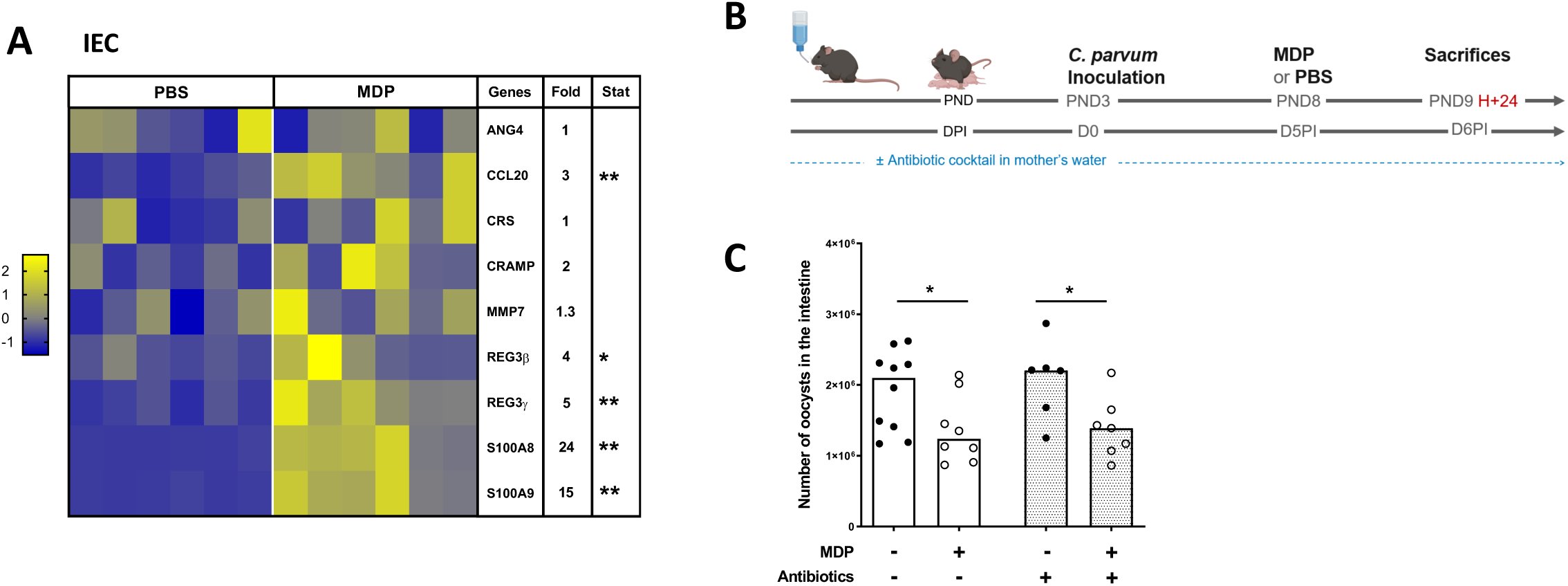
Microbiota is not involved in the MDP-induced protection despite a modified response of antimicrobial response in neonatal mice. PND3 neonatal mice were orally infected with 5.10^5^ oocysts of *C. parvum* and received 200µg of MDP by IP route at 5 d.p.i.. Mice were sacrificed 4 h post-injection to evaluate the response in antimicrobial peptides of the purified IECs by RT-qPCR (**A**). Heat map is designed by z-score of the 2^-Δct^ results for each gene and fold change of mRNA expression calculated as 2^-Δct^ results of the MDP-group in comparison to the PBS-group (n = 6 mice for each group). (**B**) Experimental timeline. Pregnant mice received a mixture of antibiotics in their drinking water 2 days before the expected day of delivery and throughout the experiment to reduce their gut intestinal bacterial burden and that of their pups. (**C**) Parasite load in the intestine was evaluated 24 h later (n = 7–10 mice for each group). Each point corresponds to one mouse and bars represent the median. Statistics are calculated by the Mann-Whitney test. ns>0.05, *P<0.05, **P<0,01. PND = postnatal days; DPI=days post-inoculation.

Among IECs, Lgr5+ stem cells constitutively express NOD2 receptor at a higher level compared to Paneth cells and other epithelial cells, contributing to their protection and epithelial regeneration (20, 23, 33). In neonatal mice infected with *C. parvum* that received MDP, a significant hyperproliferation of intestinal stem cells was observed after Ki67+ staining (Fig 5A). We next isolated small intestinal crypts from neonatal mice injected with PBS or MDP and cultured them in Matrigel^®^ to allow intestinal organoids development (Fig 5B). The organoids were allowed to mature *in vitro* for 6 days before being counted and characterized. Following *in vivo* stimulation of mice with MDP, a significant increase in the number of organoids was observed (Fig 5C), and there was an increased proportion of budded organoids, indicating enhanced maturation and regeneration of the epithelium (Fig 5D). Furthermore, the size of budded organoids increased when derived from intestinal crypts isolated from MDP-injected neonatal mice (Fig 5E). Taken together, these data clearly demonstrate that injection of MDP into *C. parvum*-infected neonatal mice stimulated, either directly or indirectly, stem cell proliferation and promoted epithelium regeneration, which could explain elimination of infected cells, and therefore the reduction in parasite numbers.

**Fig 5.**
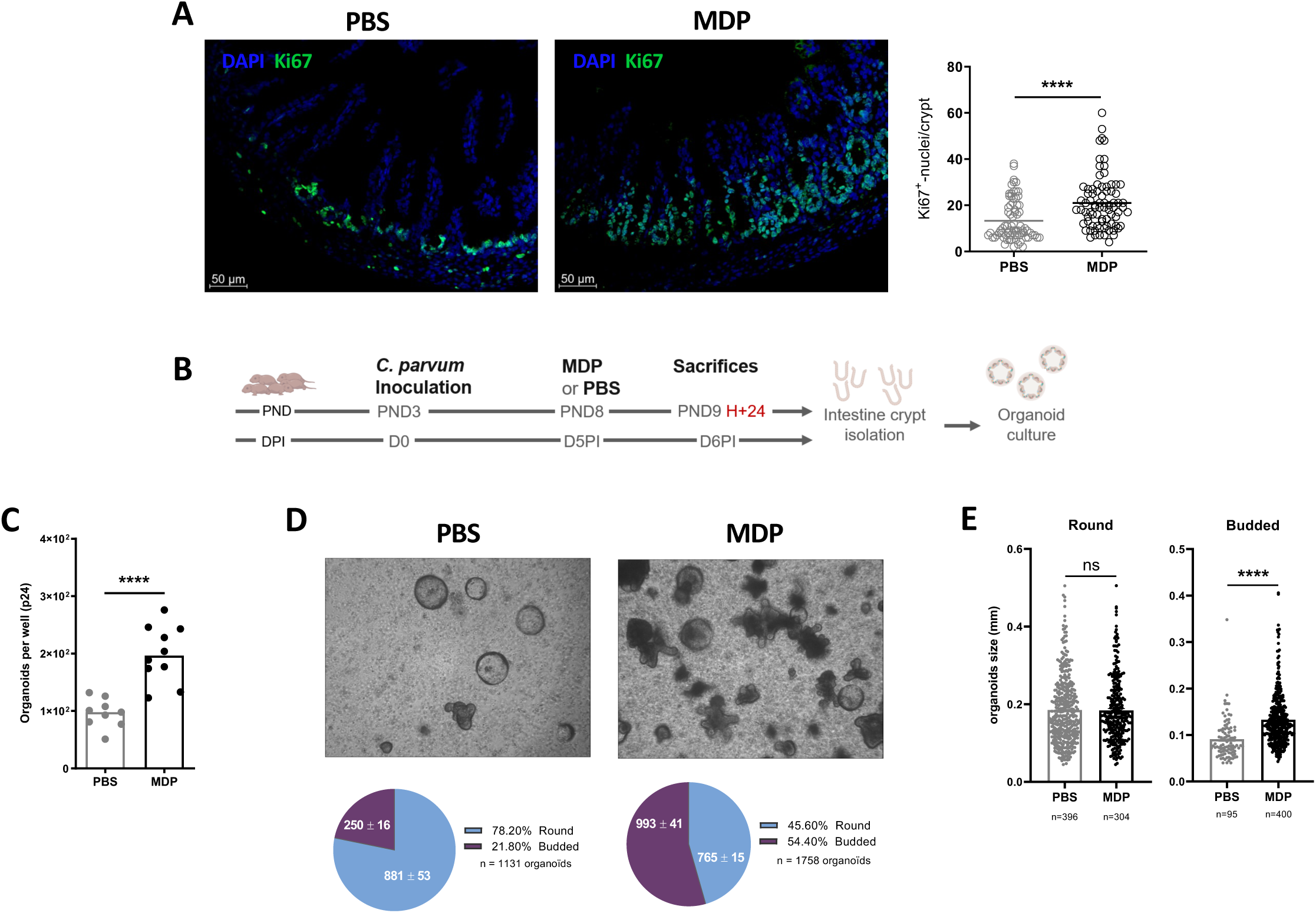
MDP injection promotes intestinal epithelial regeneration in neonatal mice infected by *C. parvum*. PND3 neonatal mice were orally infected with 5.10^5^ oocysts of *C. parvum* and received 200µg of MDP by IP route at 5 d.p.i.. Mice were scarified 24 h post-injection to evaluate the impact on IECs. (**A**) The staining of Ki67+ cells in the ileum of infected mice with or without MDP treatment was quantified by the number of Ki67+ cells per crypt in three separate areas for each animal (n = 3 animal/condition) (right panel). In the left panel, immunofluorescence staining of neonatal mice 40x, with Ki67+ cells (green) and nuclei (DAPI, blue). (**B**) Experimental timeline to generate organoids. Intestinal crypts were isolated from MDP- or PBS-infected mice (n = 5 per group) to generate and culture intestinal organoids for 6 days (2 wells of a 24-well per animal). The numbers (**C**), the state of development (**D**) and the size of organoids (**E**) were determined. PND = postnatal days. DPI = days post-inoculation. Statistics are calculated by the Mann-Whitney test. ns>0.05, ****P<0,0001.

### IFN-γ is a key cytokine involved in the mechanism underlying the protection induced by MDP in neonatal mice

IFN-γ is a key cytokine essential for host resistance to *C. parvum* (34, 35). Furthermore, it actively contributes to the reduction of parasite load in the intestine of neonatal mice upon stimulation with TLR ligands (11, 12). Therefore, we injected MDP into IFN-γ deficient neonatal mice and observed no significant reduction in parasite load compared to PBS-IFN-γ deficient neonatal mice, in contrast to what was observed in WT neonates (Fig 6A). Furthermore, in the absence of IFN-γ, we observed neither recruitment of neutrophils in the ileum of neonatal mice after MDP injection nor modulation of the percentage of monocytes/macrophages (Fig 6B). This observation suggests a correlation between the presence of IFN-γ and the recruitment of neutrophils. When the cytokine response was analyzed under these conditions, the fold increase in gene expression of IL-1β, IL-6, IL-12p40 and CXCL2 was significantly reduced in IFN-γ deficient mice compared to wildtype (Fig 6C). This suggests that the intestinal inflammatory environment induced by MDP contributes to the reduction of *C. parvum* load.

**Fig 6.**
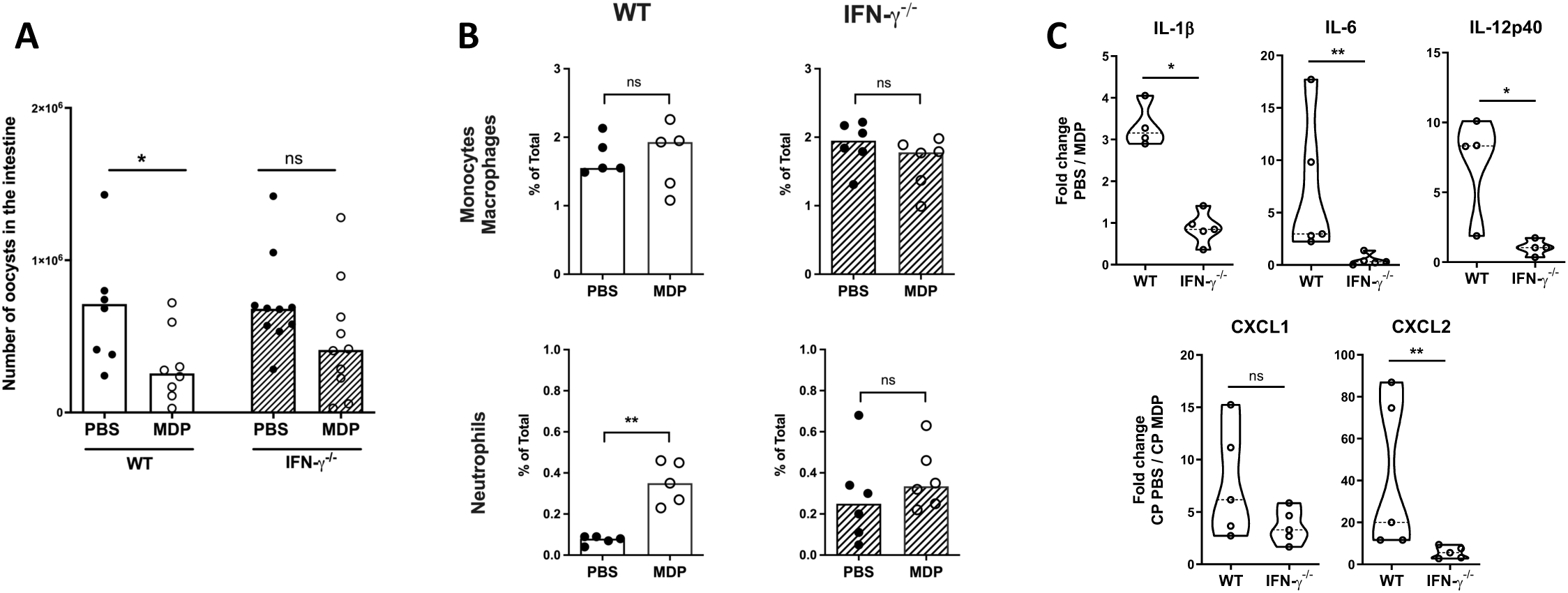
The reduced *C. parvum* intestinal load induced by MDP into neonatal mice is dependent on IFN-γ. PND3 WT or IFN-γ^-/-^ neonatal mice were orally infected with 5.10^5^ oocysts of *C. parvum* and received 200µg of MDP by IP route at 5 d.p.i.. (**A**) The intestinal parasite load in the neonatal mice was evaluated 24 h after MDP injection (n = 7–10 mice for each group). (**B-C**) Four hours after MDP-injection, intestines were collected and analyzed by flow cytometry and by qRT-PCR. (**B**) Cells from the intestinal *lamina propria* were collected and analyzed by flow cytometry. The percentage of monocytes/macrophages (CD45+ CD11c+ MHCII+ CD64+) and neutrophils (CD45+ Ly6G+ CD11b+) was determined (n = 5-6 mice for each group). (**C**) Levels of inflammatory gene expression were quantified by RT-qPCR in the ileum. Data are expressed by the fold change of mRNA expression calculated as 2^-Δct^ results of MDP-group in comparison to the PBS-group of each mouse strain (WT or IFN-γ^-/-^) (n = 4-5 mice for each group). Each point corresponds to one mouse and bars represent the median. Statistics are calculated by the Mann-Whitney test. ns>0.05, *P<0.05,**P<0,01. PND = postnatal days.

### The IFN-γ-dependent production of the cytokine IL-22 contributes to the reduction of the intestinal parasite load induced by MDP injection

The increased response of Reg-3β/γ and S100A8/A9 in IECs and the observed epithelial regeneration in the small intestine of neonatal mice following MDP injection (Fig 4A) prompted us to investigate the role of IL-22 in this process. In fact, it is well documented that IL-22 induces the production of innate antimicrobial molecules including defensins, Reg family molecules, and S100 proteins (36–39) and stimulates the wound healing of intestinal tissue (40–43). We first examined the mRNA response of IL-22 in ileal tissue and observed an approximately 10-fold increase after MDP injection (Fig 7A). This intestinal increased of IL-22 mRNA gene expression post-MDP injection was not observed in absence of IFN-γ (Fig 7A) suggesting its involvement in the IFN-γ-dependent protective mechanism induced by MDP. Consistently, the increased Reg-3β/γ and S100A8 mRNA expression was also diminished in IFN-γ deficient mice after MDP injection compared to WT neonatal mice (Fig 7A). Since IECs are the only cells in the intestine that express the IL-22 receptor (36, 44), and that also express antimicrobial peptides after MDP injection in infected neonatal mice, as shown in Fig 4A, we further investigated a possible direct protective effect of IL-22 on infected epithelial cells as has been described for other cytokines such as IFN-γ, TNF-α, and type I IFN (45–47). We then analyzed the role of IL-22 *in vivo* by injecting a neutralizing antibody against IL-22 and observed a loss of protection induced by MDP injection (Fig 7B).

**Fig 7.**
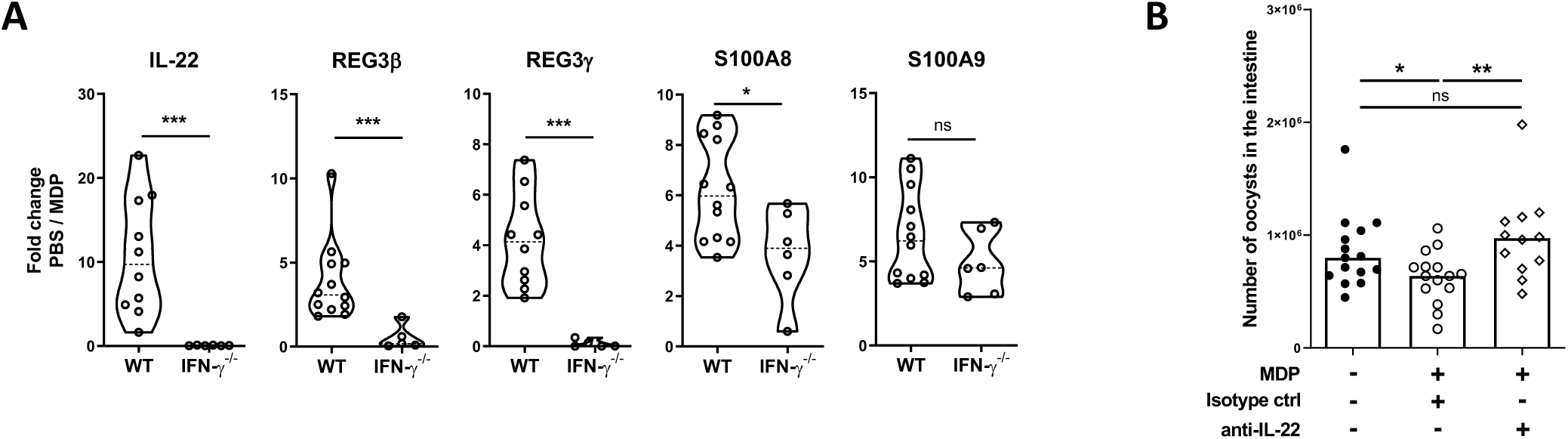
IL-22 plays a role in the protection induced by MDP in *C. parvum*-infected neonatal mice. **(A)** PND3 WT or IFN-γ^-/-^ neonatal mice were orally infected with 5.10^5^ oocysts of *C. parvum* and received 200µg of MDP by IP route at 5 d.p.i.. Mice were sacrificed 4 h post-injection to evaluate the intestinal (ileal tissue) gene expression levels of IL-22, REG3β/γ, S100A8/A9 by RT-qPCR. Data are expressed by the fold change of mRNA expression calculated as 2^-Δct^ results of MDP-group in comparison to the PBS-group of each mouse strain (WT or IFN-γ^-/-^) (n = 4-5 mice for each group). (**B**) PND3 WT neonatal mice were orally infected with 5.10^5^ oocysts of *C. parvum* and mice received 200µg of MDP and an anti-IL-22-neutralizing antibody or isotype control antibody by IP route at 5 d.p.i.. Mice were sacrificed 24H post-injection to evaluate the parasite load in the intestine (n = 12–15 mice for each group). Each point corresponds to one mouse and bars represent the median. Statistics are calculated by the Mann-Whitney test. ns>0.05, *P<0.05,**P<0,01, ***P<0,001. PND = postnatal days.

Altogether, these data demonstrate that IFN-γ contributes to an immune loop involving neutrophils, IL-22, Reg3β/γ and S100A8/A9 that helps to reduce *C. parvum* intestinal load after MDP injection.

## Discussion

Early-life is characterized by increased susceptibility to respiratory and enteric infections. In humans, neonatal infections remain a major concern, with 5 million children under the age of 5 succumbing from infectious diseases in 2021 (United Nations Inter-Agency Group for Child Mortality Estimation (UN IGME), 2023). Similarly, respiratory and intestinal infections in calves and piglets result in significant economic losses in livestock production. High morbidity rates have been reported, such as diarrhea (calves ≤ 35%; piglets ≤ 50%) and respiratory diseases (calves ≤ 80%; piglets ≤ 40%)(48). This vulnerable period is characterized by quantitative and qualitative defects in immunity, particularly at mucosal surfaces, where environmental, nutritional, and microbial antigens shape the immune system. Newborns, lacking a fully developed adaptive immune system, rely on innate immune responses and maternally transmitted immunity for early protection against pathogens (49). Enhancing protection against infection and disease by modulating the innate immune response is therefore a feasible approach.

Modulation of Toll-like receptors and associated pathways presents opportunities to improve pathogen recognition and modulate the inflammatory response to combat infection. As an example, prophylactic TLR2 activation can prime airway immunity for enhanced protection against rhinovirus, influenza and SARS-CoV-2 infection (50, 51). In addition, PolyI:C treatment to stimulate TLR3 has been described to effectively promote the clearance of hepatitis B virus (HBV) infection (52). In neonates, the benefits of administering TLR agonists (e.g., resiquimod and LPS) have been demonstrated in a neonatal mouse model predisposed to poor sepsis outcomes (53). In addition, we have previously demonstrated the benefits of TLR9 and TLR3 stimulation by administration of CpG-ODN and PolyI:C respectively to induce a strong intestinal innate immune response and efficiently control cryptosporidiosis in the neonatal mouse model (11, 12).

In this study, we have demonstrated that MDP, the agonist of the NOD2 receptor in neonatal mice can stimulate both systemic and intestinal inflammatory responses, resulting in rapid renewal of the intestinal epithelium, which transiently helps to clear *C. parvum* from the intestine. Mechanistically, mice were protected upon MDP injection in an IFN-γ and IL22-dependent manner. Similarly, NOD2 activation by bacterial MDP has previously been described to suppress viral infection by human cytomegalovirus (HCMV) via an IFN-β-dependent pathway (54) and to protect from oxazolone-induced colitis by stimulating hematopoietic cells (55). Our previous studies, using TLR3 and TLR9 ligands to stimulate the innate immune system of neonatal mice and protect them against cryptosporidiosis, highlighted the crucial role of MyD88 signaling activation in dendritic cells. When TLR3-TRIF signaling pathway is activated by PolyI:C, the presence of microbiota is required to allow the simultaneous MyD88-activation via a TLR5 recognition (12). In this study, the antibiotic-induced microbiota modifications had no effect in the MDP-induced protection, suggesting that activation of the NOD2 signaling pathway is sufficient to promote parasite clearance.

In the present study, the reduced infection of *C. parvum* induced in neonatal mice by MDP injection was dependent on IFN-γ and its deficiency was associated with a diminished recruitment of neutrophils and an altered response in IL-22 and Reg3γ. To allow MDP to be delivered to the intracellular receptor NOD2, the di-tripeptide transporter PepT1 plays a critical role (56, 57). Interestingly, the expression of the transporter (58) is resistant to intestinal mucosal injury (59–62) and is increased by IFN-γ (63) and during Cryptosporidiosis in a rat model (64). Taken together, these data may give new insights in the IFN-γ dependent mechanism of protection induced by MDP administration we described in this study.

We demonstrated that NOD2 stimulation in neonates promoted a rapid influx of neutrophils into the intestine attributable to the increased CXCL1 and CXCL2 response of IECs, whereas the proportion of macrophages remains unchanged. Neutrophils are recruited into the intestine after *C. parvum* infection (7, 65, 66) but their role in the control of cryptosporidiosis is still poorly understood. In an *in vitro* study, neutrophils were shown to form NETs that resulted in the entrapment of 15% of sporozoites (67). However, *in vivo* depletion of neutrophils in the piglet did not affect the severity or pathology of infection (66). In the SCIDbgMN mouse model, which is characterized by a depletion of functional macrophages and neutrophils in a SCIDbg background already lacking T, B, and NK cells, Takeuchi *et al.* reported a more severe parasite load in the early phase of *C. parvum* infection, leading to mouse mortality (68). Neutrophils have been proposed to convert resident “M2” macrophages to an “M1” macrophage phenotype, contributing to the mounting of a protective immune response (68). Their role in cryptosporidiosis may therefore be less direct and more in cooperation with other cell populations. In the present study, when neutrophils and macrophages were sorted to analyze their transcriptional profile, both cell types exhibited higher levels of IL-1β and IL-6 after MDP injection, and macrophages decreased their TGF-β and IL-13 mRNA response. These data may be related to a potential conversion of macrophages toward a pro-inflammatory profile associated with the neutrophil influx in the intestine of neonates after NOD2 stimulation, which may help to reduce *C. parvum* infection.

We also reported a robust IFN-γ-dependent production of IL-22 after MDP injection in the intestine of neonates, which contributed to the reduction of the parasite burden. Surprisingly, IL-22 expression in neutrophils of neonatal mice was not significantly modified by MDP injection (S4 Fig.). However, consistent with the lack of neutrophil recruitment in IFN-γ deficient mice, it could be suggested that the increased intestinal IL-22 response is due to the increased number of neutrophils. Several studies have described that IL-22 induces the production of antimicrobial peptides such as Reg3γ and is involved in the recovery of the intestinal epithelial barrier after acute injury by promoting the proliferation of LGR5+ intestinal epithelial stem cells. In addition, IL-22-mediated regulation of epithelial function closely parallels that of other pro-inflammatory cytokines, such as IFN-γ and TNF-α (69). We observed a significant increase in the Reg3γ mRNA after MDP injection, which was reduced in the absence of IFN-γ, where the IL-22 response was also significantly reduced. In addition, we described that the systemic injection of MDP in neonates affected intestinal epithelial stem cell proliferation. Taken together these data suggest that the classical interplays involved in the maintenance of intestinal homeostasis between IFN-γ, IL-22, Reg3γ and the intestinal epithelium, would allow the elimination of infected cells in *C. parvum* infected neonatal mice after MDP injection.

In this work, we have shown for the first time, to the best of our knowledge, that stimulation of the NOD2 receptor can help to clear an intestinal infection in neonatal mice. Future studies will be essential to investigate the precise role of IL-22 in the protection induced by MDP to combat cryptosporidiosis and to maintain an efficient protective immune response.

## Materials and Methods

### Ethic statements

All experimental protocols were performed in accordance with French legislation (Décret: 2001-464 29/05/01) and EEC regulations (86/609/CEE) on the care and use of laboratory animals, after validation by the local ethics committee for animal experimentation (Comité d’Ethique pour l’Expérimentation Animale Val de Loire (CEEA VdL n°019: APAFIS#34587, APAFIS#21515). The mice were housed in a specific pathogen-free animal facility in the Infectiology of Farm, Model and Wildlife Animals Facility (PFIE, Centre INRAE Val de Loire: https://www6.val-de-loire.inrae.fr/pfie/ member of the National Infrastructure EMERG’IN). They were housed in individual ventilated cages with HEPA filters, with food and water *ad libitum*. All personnel involved in animal work received specific training in animal care, handling and experimentation, as required by the French Ministry of Agriculture.

### Parasite preparation

Oocysts of *C. parvum* were originally obtained from an infected child (INRAE *C. parvum*). This strain was continuously maintained by regular passages in newborn calves at the PFIE Centre INRAE Val de Loire facility and oocysts were purified following a protocol previously described (34). Transgenic INRAE *C. parvum* expressing Nano-luciferase (70) was used, for *in vitro* experiments. This strain was maintained by regular passages in IFN-γ^-/-^ mice.

### Mouse infection and experimental models

Specific-pathogen-free C57BL/6J mice (WT and Interferon-gamma knock-out (IFN-γ^-/-^)) were used in this study. Three-day-old neonatal mice (PND3 for postnatal day) were orally inoculated with 5.10^5^ oocysts of *C. parvum*. Five days post-inoculation (d.p.i.), neonatal mice received NOD ligands or PBS, and parasitic load was determined 24 hours (h) later. The level of infection in individual neonatal mice was assessed by counting the number of oocysts in the whole intestine, as previously described (71). NOD1 agonist, iEDAP: 200µg (tlrl-dap, Invivogen), NOD2 agonist, MDP: 5µg, 50µg, 200µg (tlrl-mdp, Invivogen), MDP control: 200µg (MDP Ct) (tlrl-mdpcl, Invivogen) were used in this study. To decipher the innate immune response, infected mice were sacrificed 4 h or 24 h after NOD ligand administration and different samples were collected for further analysis: ileum, spleen and peritoneal cells. To study the impact of microbiota in the MDP-induced-protection mechanism, dams of newborns were treated with an antibiotic solution dissolved in drinking water from two days before delivery until analysis of the neonates at 6 d.p.i.. The antibiotic solution was composed of ampicillin (1mg/ml), vancomycin (0.5mg/ml), colistin (1mg/ml), streptomycin (5mg/ml) (Antibiotics, Sigma-Aldrich) and 2.5% (wt/vol) sucrose. To evaluate the role of IL-22, infected mice received simultaneous intraperitoneal injections of 200µg of MDP and 10µg of mouse IL-22 antibody (R&D Systems) or an equivalent amount of isotype IgG from goat (R&D Systems), while control mice received PBS.

### Immune Cell preparation

Peritoneal cells were obtained from the peritoneal lavage of PND8 C57BL/6 mice. Peritoneal cells were collected by injecting 4 x 500 µl of RPMI into the peritoneal cavity. The cells were then passed through a 100 μm strainer and centrifuged. Finally, the cells were distributed in round-bottom 96-well plates (Falcon) for flow cytometry analyses or were frozen at -80°C with TRI Reagent (Sigma-Aldrich) for future transcriptomic studies. To isolate immune cells from *lamina propria*, jejunum and ileum were cut longitudinally, washed with RPMI medium, placed in ice-cold Cell Recovery Solution (Corning) and incubated for 3 hours at 4°C. The intestines were then vigorously shaken to separate the IECs from the *lamina propria*. The contents were filtered through a 100 µm cell strainer to isolate the IECs. The IECs was centrifuged at 400g for 10 min at 4°C and the pellets were frozen at -80°C with TRI Reagent (Sigma-Aldrich) and preserved for future transcriptomic studies. The intestines retained on the filters were mechanically cut into 0.5 cm pieces and enzymatically digested in RPMI 10% FCS with 100U/mL of collagenase type I (Sigma-Aldrich) for 1h at 37°C. After digestion, cells were filtered through 200 µm filters and centrifuged at 400g for 10 min at 4°C. Pellets were washed with 10% FCS in PBS and cells were collected with PBS 2% FCS 1% mouse serum to block non-specific staining. Finally, *lamina propria* cells were distributed in round-bottom 96-well plate (Falcon) for flow cytometry analysis or cell sorting.

### Flow cytometry and cell sorting

Isolated cells were stained with a mix of antibodies in a FACS medium (PBS, 2% FCS, 2 mM EDTA) for 30 min in the dark at 4°C. The mix contained anti-mouse CD45-BV711 (BioLegend); MHCII-BV421 (BioLegend); CD11c-FITC (BioLegend); CD64-PE (Miltenyi Biotec); CD11b-APC (BioLegend); Ly6G-PeCy7 (BioLegend). Cells were then washed twice then fixed with fixation buffer (BD Biosciences) for flow cytometry, or suspended in PBS, 2% FCS, 2 mM EDTA for cell sorting. Cells were analyzed on an LSR Fortessa X-20 (BD Biosciences) or sorted using an high speed cell sorter MoFlo Astrios EQ (Beckman Coulter). Flow cytometry data were analyzed using Kaluza software (Beckman Coulter). Sorted cells were stored for future transcriptomic studies. S1 Fig shows the gating strategy used for the analyses.

### RNA extraction and qRT-PCR

At sacrifice, spleen and ileum tissues were frozen in nitrogen liquid and stored at -80°C. Total RNAs (spleen, ileum, peritoneal cells) were extracted using Trizol reagent solution (Sigma-Aldrich) and processed according to the manufacturer’s recommendations. Reverse transcriptase (RT) reaction was performed using oligo(dT)18 primers and random hexamers with M-MLV reverse transcriptase (Promega). RT-qPCR reaction consisted of 7.5µL of SybrGreen (BIO-RAD), 0.5µM of forward primer, 0.5µM of reverse primer, and RNase DNase-free water up to 13µL. Finally, 20ng of cDNA samples were added. Quantitative RT-PCR was performed on a CFX96 instrument (Biorad) and the cycling steps were 5 min at 95°C, followed by 40 cycles of 10 s at 95°C, 15 s at 60°C, and a melting curve phase. Expression levels were calculated using the formula 2^-ΔCt^, where ΔCt=Ct gene−Ct average housekeeping for the control sample. HPRT and PPia were used as internal standards for normalization. Primers for gene quantification are described in S2 Fig. Heat maps are constructed using the z-score of the 2^-ΔCt.^

### Immunofluorescence imaging and analysis

Samples of ileum tissues were fixed overnight at 4°C in a fresh solution of 4% para formaldehyde (Sigma-Aldrich) in PBS. Samples were then washed for 1 day in PBS and incubated in a solution of 30% sucrose-PBS (Sigma-Aldrich) for at least 3 hours. Ilea were included in paraffin blocks and sliced. Sections were de-waxed by an automate Leica ST5020 Multistainer (Leica biosystem) and incubated in a sodium citrate buffer (pH = 6.12) for 20 min for antigen retrieval. Sections were next permeabilized and Fc-receptors were blocked using a solution of PBS, 0,1% TritonX-100, 10% BSA for 20 min at room temperature (RT). Sections were stained for 1h at RT using a primary antibody rabbit anti-Ki67 (Abcam) at 1/100 in PBS, 0,1% Triton, 1% BSA. After washing, a secondary antibody, swine anti-rabbit FITC (DAKO) was added at 1/100 in PBS, 0,1% Triton, 1% BSA for 1h at RT. Cell nuclei were counterstained with DAPI 1/2000 for 10 min at RT. Slides were mounted using Fluoromount-G medium (Invitrogen). The acquisitions were performed under a widefield fluorescence microscope (Zeiss Axiovert 200M, Zeiss). Acquisitions were realized using a 40x objective using Zen Imaging software (Zeiss). Images were taken from 3 separate areas for each animal and blindly analyzed with the ImageJ software (version 2.9.0/1.53t, https://imagej.nih.gov). A semi-automated homemade macro was used to quantify the number of Ki67^+^-nuclei per crypt. Briefly, crypts were manually segmented. Crypts were individually cropped, and StarDist (72, 73) (version 0.8.3, https://github.com/stardist) was used to segment nuclei based on DAPI signal. By applying a threshold to the Ki67 signal, the number of proliferative cells per crypt was determined and compared between conditions.

### Isolation of intestinal crypts and organoid culture

Sections of ileal segments were removed from infected mice that received or not MDP and placed in PBS without Ca^2+^ and Mg^2+^ (ThermoFisher Scientific) supplemented with 100 U/mL penicillin and 100 mg/mL streptomycin. Intestinal crypts were isolated according to established protocols described previously (74, 75). The tissue was opened longitudinally and cut into pieces. These pieces were then immersed in a dissociation buffer consisting of PBS without Ca2+ and Mg2+ containing 9 mM EDTA (Sigma Aldrich), 3 mM 1,4-Dithiothreitol, (DTT, Sigma Aldrich) and 10 µM Y27632 (Tocris) to dissociate intestinal crypts during 45 minutes of continuous shaking at 16 rpm at RT. The intestinal fragments were then transferred to a cold PBS solution without Ca2+ and Mg2+ and gently shaken by hand in an up-and-down motion for 2 minutes. To eliminate the intestinal villi, the supernatant was first passed through a 100 µm cell strainer and then rinsed with PBS without Ca2+ and Mg2+. This process was repeated using a 70 µm cell strainer and the crypts were then centrifuged at 220g for 5 minutes at RT. The pellet containing intestinal crypts was suspended in DMEM/F12-Glutamax (Gibco Life Technologies), 1% HEPES (Gibco Life Technologies), 5% FBS, 100 U/mL penicillin and 100 mg/mL streptomycin. Approximately 2500 crypts were seeded into 50µl of 75% of Matrigel™ mixed with 25% complete medium (DMEM/F-12 Glutamax (Gibco Life Technologies) supplemented with 50% L-WRN CM (homemade as described before (75)), 10 mM HEPES (Gibco Life Technologies), B27 1X (ThermoFisher Scientific), 50 ng/mL EGF (Sigma Aldrich), 500 nM A83-01 (Tocris), 10 µM SB2022190 (Tocris), 10 nM gastrin I (Tocris), 1 mM *N*-acetyl-L-cysteine (Sigma) and 10 µM Y27632 (Tocris)) and seeded into pre-warmed 24-well plates. After Matrigel™ polymerization, 500 μL of complete medium was added per well. The organoid medium was refreshed every 2-3 days with complete medium. Images were captured by inverted phase contrast microscopy (Olympus CK40) and with an OptikaB9 digital camera (Optika). The number, size and development stage of the organoids were monitored using the Optikavision Lite 2.1 software.

### Statistics

Statistical analyses were performed using the GraphPad Prism v8 software (GraphPad Software, San Diego, USA). The non-parametric Mann-Whitney t-test was used to determine the significance between two groups. For comparisons involving more than two groups, the one-way non-parametric test (Kruskall-Wallis) was used. P-values less than 0.05 were considered statistically significant.

## Acknowledgments

We extend our gratitude to Nathalie Lallier for her assistance in mouse experimentation, Emilie Doz-Deblauwe for discussions on neutrophils, Christelle Rossignol for providing expert guidance and technical support in histology. We are also grateful to Valérie Quesniaux and Jean-Marc Cavaillon for their valuable discussions and critical expertise, which greatly contributed to advancing our reflections. Additionally, we acknowledge the valuable contributions of Fanny Faurie, Thierry Chaumeil, Emilie Lortscher, and Corinne Beaugé at INRAE-PFIE for rearing and supplying mice. All experimental timelines were created using BioRender.com. The English editing was checked using the Deep Writing software.

## Author Contributions

**Conceptualization**: Sonia Lacroix-Lamandé

**Data Curation**: Mégane Fernandez, Tiffany Pezier, Sonia Lacroix-Lamandé

**Formal analysis:** Mégane Fernandez, Tiffany Pezier, Julien Pichon, Sonia Lacroix-Lamandé

**Funding acquisition:** Sonia Lacroix-Lamandé, Catherine Werts

**Investigation:** Mégane Fernandez, Tiffany Pezier, Sonia Lacroix-Lamandé

**Methodology:** Mégane Fernandez, Tiffany Pezier, Sonia Lacroix-Lamandé

**Project administration:** Sonia Lacroix-Lamandé

**Resources:** Mégane Fernandez, Tiffany Pezier, Yves Le Vern, Sonia Lacroix-Lamandé

**Supervision:** Sonia Lacroix-Lamandé

**Validation:** Mégane Fernandez, Tiffany Pezier, Sonia Lacroix-Lamandé

**Visualization:** Mégane Fernandez, Sonia Lacroix-Lamandé

**Writing – original draft:** Mégane Fernandez, Sonia Lacroix-Lamandé

**Writing – review & editing:** Mégane Fernandez, Julien Pichon, Yves Le Vern, Catherine Werts, Sonia Lacroix-Lamandé

## Supporting information captions

**S1 Fig.**
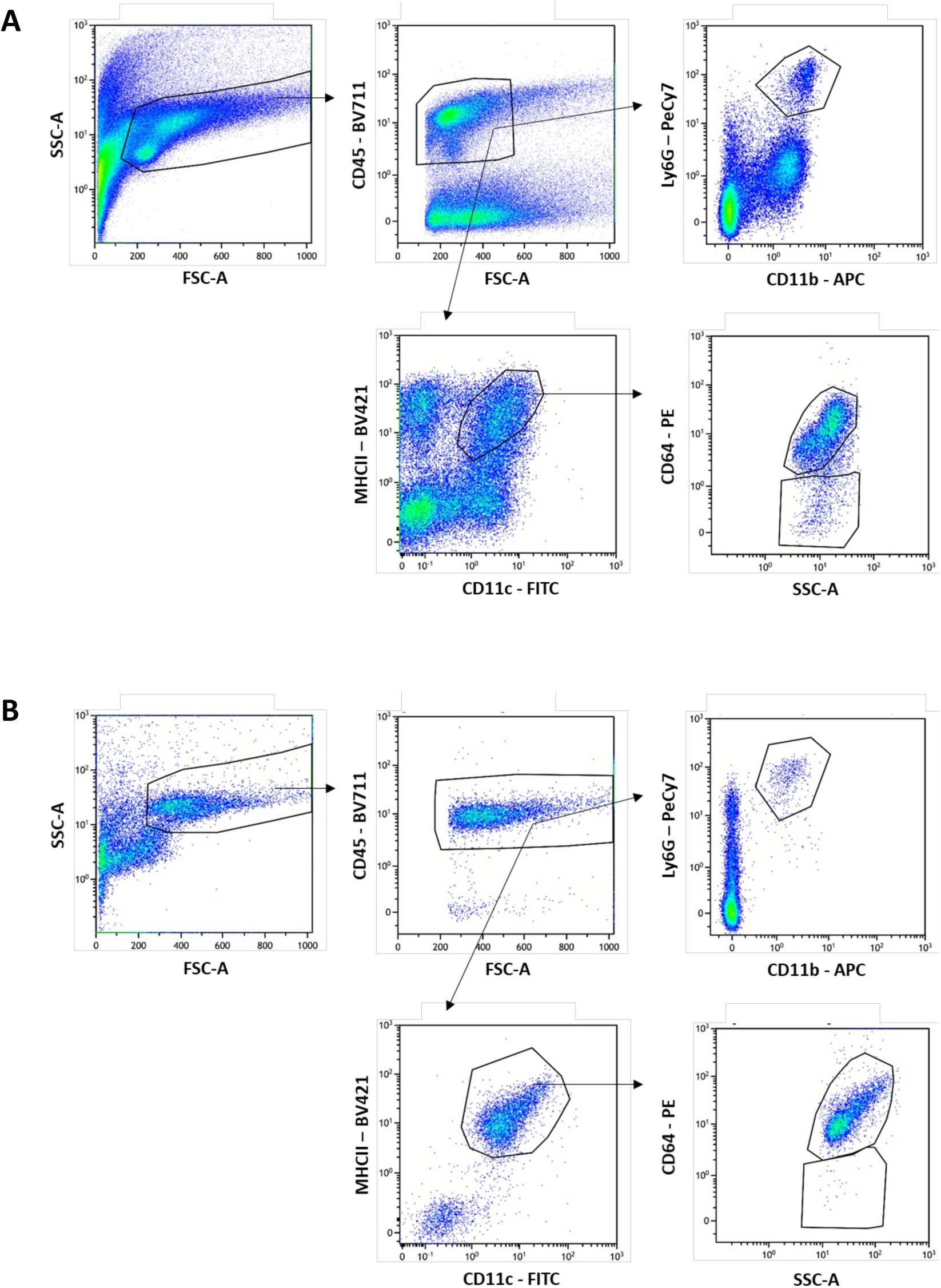
Gating strategies for multicolor flow cytometry on lamina propria cells (A) or on IP cells (B). Neutrophils: CD45+ Ly6G+ CD11b+. Monocytes/Macrophages: CD45+ CMHII+ CD11c+ CD64+. Dendritic cells: CD45+ CMHII+ CD11c+ CD64-

**S2 Fig.**
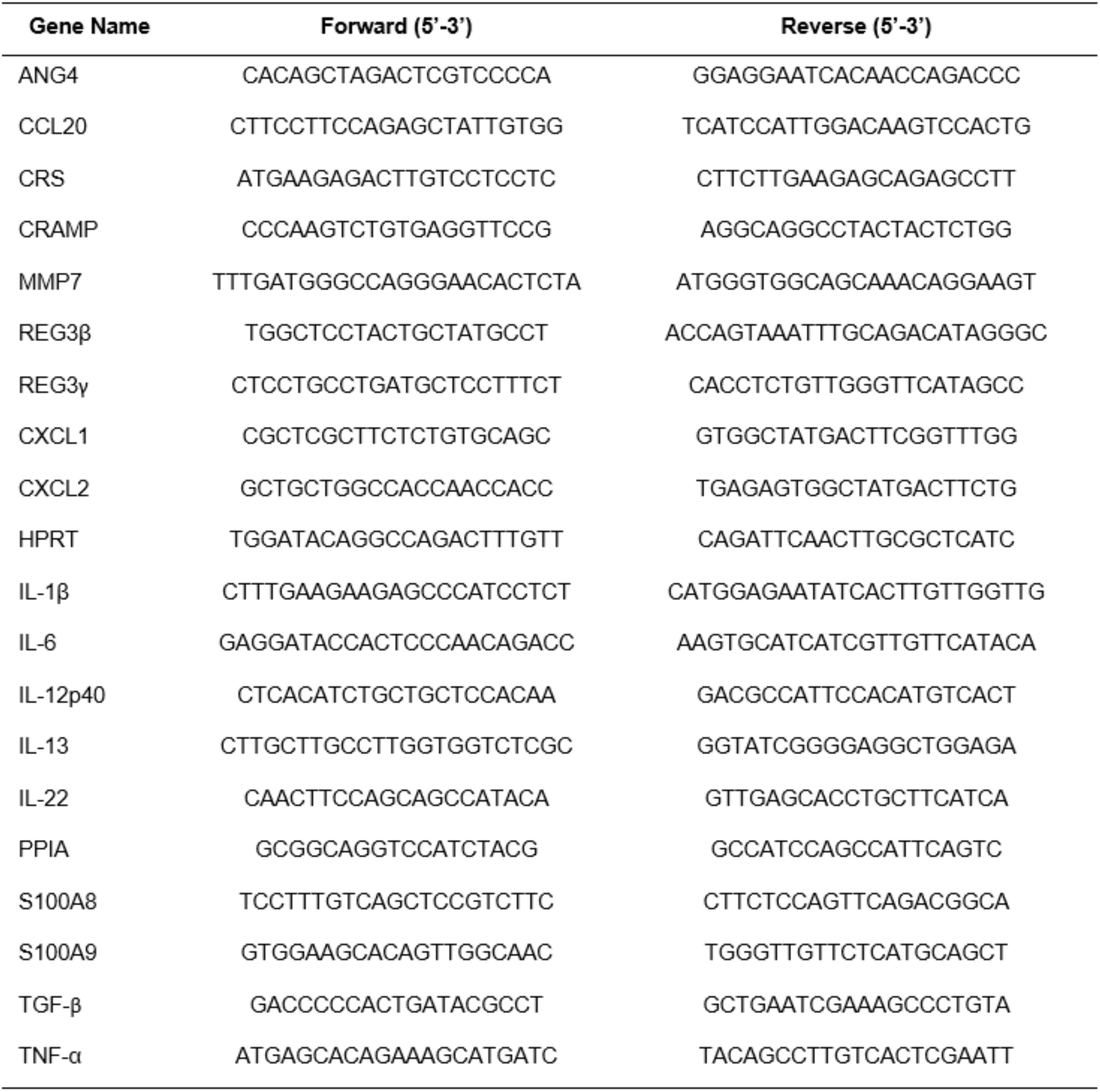
**Primer sequences used for q-RT-PCR**

**S3 Fig.**
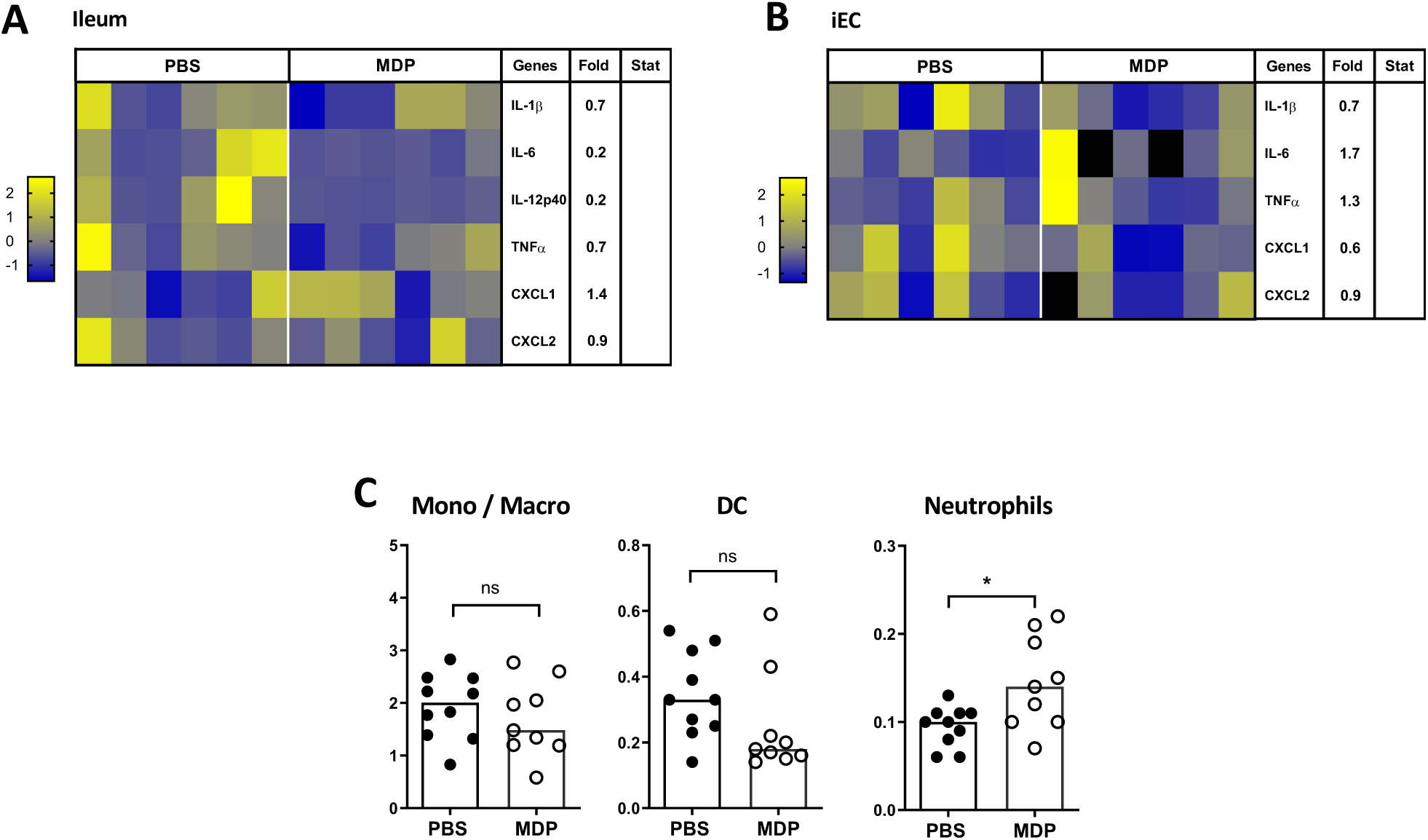
MDP injection has an impact on neutrophils recruitment in the intestine of neonatal mice infected by *C. parvum* 24 h later PI. Neonatal mice were orally infected with 5.10^5^ oocysts of *C. parvum* at 3-days-old and received an injection of MDP by IP route (200µg) 5 d.p.i.. 24 h after MDP injection, the intestine was sampled. (**A-B**) The levels of inflammatory gene expression were quantified by RT-qPCR in the ileum tissue (**A**) and in purified intestinal epithelial cell (IECs) (**B**). Heat maps are designed by z-score of the 2^-Δct^ results for each gene and fold change of mRNA expression calculated as 2^-Δct^ results of the MDP-group in comparison to the PBS-group (n = 6 mice for each group). (**C**) Cells from the intestinal *lamina propria* were collected and analyzed by flow cytometry. The percentage of monocytes/macrophages CD45+ CD11c+ MHCII+ CD64+, dendritic cells (DC) CD45+ CD11c+ MHCII+ CD64-, neutrophils CD45+ Ly6G+ CD11b+ was determined. (n = 9-10 mice for each group). Statistics are calculated by the Mann-Whitney test. ns >0.05, *P<0.05.

**S4 Fig.**
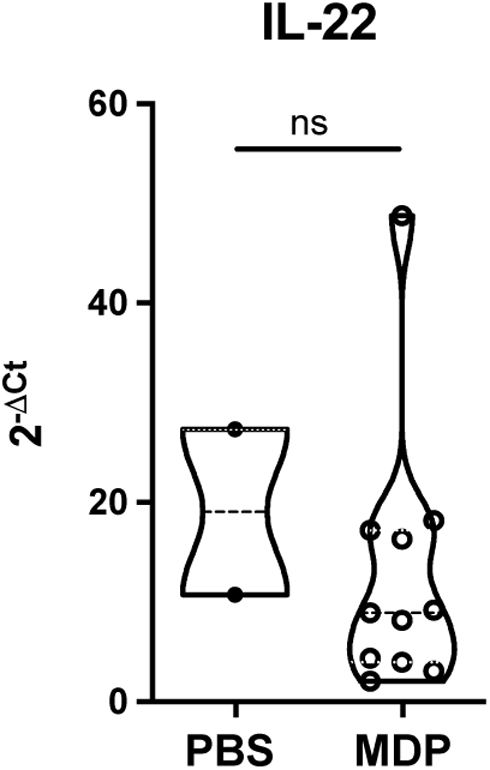
MDP injection does not change IL-22 expression in neutrophils from *C. parvum*-infected neonatal mice. PND3 neonatal mice were orally infected with 5.10^5^ oocysts of *C. parvum* and received 200µg of MDP by IP route at day 5 d.p.i.. 4 h after MDP injection, the intestine was sampled. Cells from the *lamina propria* of the distal small intestine were collected. Neutrophils were sorted, and the levels of IL-22 expression was quantified by RT-qPCR. Results are expressed as the 2^-Δct^ compared between the MDP-group (n = 11 mice) and the PBS-group (n = 2, pool of 9 mice). Statistics are calculated by the Mann-Whitney test. ns>0.05.

## References

1. Thomson S, Hamilton CA, Hope JC, Katzer F, Mabbott NA, Morrison LJ, et al. Bovine cryptosporidiosis: impact, host-parasite interaction and control strategies. Vet Res. 2017;48(1):42.

2. Innes EA, Chalmers RM, Wells B, Pawlowic MC. A One Health Approach to Tackle Cryptosporidiosis. Trends Parasitol. 2020;36(3):290–303.

3. Striepen B. Parasitic infections: Time to tackle cryptosporidiosis. Nature. 2013;503(7475):189–91.

4. Lantier L, Lacroix-Lamande S, Potiron L, Metton C, Drouet F, Guesdon W, et al. Intestinal CD103+ dendritic cells are key players in the innate immune control of Cryptosporidium parvum infection in neonatal mice. PLoS Pathog. 2013;9(12):e1003801.

5. Korbel DS, Barakat FM, Di Santo JP, McDonald V. CD4+ T cells are not essential for control of early acute Cryptosporidium parvum infection in neonatal mice. Infect Immun. 2011;79(4):1647–53.

6. Laurent F, Lacroix-Lamande S. Innate immune responses play a key role in controlling infection of the intestinal epithelium by Cryptosporidium. Int J Parasitol. 2017;47(12):711–21.

7. Laurent F, Eckmann L, Savidge TC, Morgan G, Theodos C, Naciri M, et al. Cryptosporidium parvum infection of human intestinal epithelial cells induces the polarized secretion of C-X-C chemokines. Infect Immun. 1997;65(12):5067–73.

8. Auray G, Lacroix-Lamande S, Mancassola R, Dimier-Poisson I, Laurent F. Involvement of intestinal epithelial cells in dendritic cell recruitment during C. parvum infection. Microbes Infect. 2007;9(5):574–82.

9. Potiron L, Lacroix-Lamande S, Marquis M, Levern Y, Fort G, Franceschini I, et al. Batf3-Dependent Intestinal Dendritic Cells Play a Critical Role in the Control of Cryptosporidium parvum Infection. J Infect Dis. 2019;219(6):925–35.

10. de Sablet T, Potiron L, Marquis M, Bussiere FI, Lacroix-Lamande S, Laurent F. Cryptosporidium parvum increases intestinal permeability through interaction with epithelial cells and IL-1beta and TNFalpha released by inflammatory monocytes. Cell Microbiol. 2016;18(12):1871–80.

11. Barrier M, Lacroix-Lamande S, Mancassola R, Auray G, Bernardet N, Chausse AM, et al. Oral and intraperitoneal administration of phosphorothioate oligodeoxynucleotides leads to control of Cryptosporidium parvum infection in neonatal mice. J Infect Dis. 2006;193(10):1400–7.

12. Lantier L, Drouet F, Guesdon W, Mancassola R, Metton C, Lo-Man R, et al. Poly(I:C)- induced protection of neonatal mice against intestinal Cryptosporidium parvum infection requires an additional TLR5 signal provided by the gut flora. J Infect Dis. 2014;209(3):457–67.

13. Moreira LO, Zamboni DS. NOD1 and NOD2 Signaling in Infection and Inflammation. Front Immunol. 2012;3:328.

14. Maisonneuve C, Bertholet S, Philpott DJ, De Gregorio E. Unleashing the potential of NOD- and Toll-like agonists as vaccine adjuvants. Proc Natl Acad Sci U S A. 2014;111(34):12294–9.

15. Akira S, Uematsu S, Takeuchi O. Pathogen recognition and innate immunity. Cell. 2006;124(4):783–801.

16. Ogura Y, Inohara N, Benito A, Chen FF, Yamaoka S, Nunez G. Nod2, a Nod1/Apaf-1 family member that is restricted to monocytes and activates NF-kappaB. J Biol Chem. 2001;276(7):4812–8.

17. Trindade BC, Chen GY. NOD1 and NOD2 in inflammatory and infectious diseases. Immunol Rev. 2020;297(1):139–61.

18. Hugot JP, Chamaillard M, Zouali H, Lesage S, Cezard JP, Belaiche J, et al. Association of NOD2 leucine-rich repeat variants with susceptibility to Crohn’s disease. Nature. 2001;411(6837):599–603.

19. Ogura Y, Bonen DK, Inohara N, Nicolae DL, Chen FF, Ramos R, et al. A frameshift mutation in NOD2 associated with susceptibility to Crohn’s disease. Nature. 2001;411(6837):603–6.

20. Ferrand A, Al Nabhani Z, Tapias NS, Mas E, Hugot JP, Barreau F. NOD2 Expression in Intestinal Epithelial Cells Protects Toward the Development of Inflammation and Associated Carcinogenesis. Cell Mol Gastroenterol Hepatol. 2019;7(2):357–69.

21. Penack O, Smith OM, Cunningham-Bussel A, Liu X, Rao U, Yim N, et al. NOD2 regulates hematopoietic cell function during graft-versus-host disease. J Exp Med. 2009;206(10):2101–10.

22. Ogura Y, Lala S, Xin W, Smith E, Dowds TA, Chen FF, et al. Expression of NOD2 in Paneth cells: a possible link to Crohn’s ileitis. Gut. 2003;52(11):1591–7.

23. Nigro G, Rossi R, Commere PH, Jay P, Sansonetti PJ. The cytosolic bacterial peptidoglycan sensor Nod2 affords stem cell protection and links microbes to gut epithelial regeneration. Cell Host Microbe. 2014;15(6):792–8.

24. Hisamatsu T, Suzuki M, Reinecker HC, Nadeau WJ, McCormick BA, Podolsky DK. CARD15/NOD2 functions as an antibacterial factor in human intestinal epithelial cells. Gastroenterology. 2003;124(4):993–1000.

25. Alnabhani Z, Montcuquet N, Biaggini K, Dussaillant M, Roy M, Ogier-Denis E, et al. Pseudomonas fluorescens alters the intestinal barrier function by modulating IL-1beta expression through hematopoietic NOD2 signaling. Inflamm Bowel Dis. 2015;21(3):543–55.

26. Kobayashi KS, Chamaillard M, Ogura Y, Henegariu O, Inohara N, Nunez G, et al. Nod2-dependent regulation of innate and adaptive immunity in the intestinal tract. Science. 2005;307(5710):731–4.

27. Ramanan D, Tang MS, Bowcutt R, Loke P, Cadwell K. Bacterial sensor Nod2 prevents inflammation of the small intestine by restricting the expansion of the commensal Bacteroides vulgatus. Immunity. 2014;41(2):311–24.

28. Couturier-Maillard A, Secher T, Rehman A, Normand S, De Arcangelis A, Haesler R, et al. NOD2-mediated dysbiosis predisposes mice to transmissible colitis and colorectal cancer. J Clin Invest. 2013;123(2):700–11.

29. Geddes K, Rubino S, Streutker C, Cho JH, Magalhaes JG, Le Bourhis L, et al. Nod1 and Nod2 regulation of inflammation in the Salmonella colitis model. Infect Immun. 2010;78(12):5107–15.

30. Kim YG, Kamada N, Shaw MH, Warner N, Chen GY, Franchi L, et al. The Nod2 sensor promotes intestinal pathogen eradication via the chemokine CCL2-dependent recruitment of inflammatory monocytes. Immunity. 2011;34(5):769–80.

31. Travassos LH, Carneiro LA, Ramjeet M, Hussey S, Kim YG, Magalhaes JG, et al. Nod1 and Nod2 direct autophagy by recruiting ATG16L1 to the plasma membrane at the site of bacterial entry. Nat Immunol. 2010;11(1):55–62.

32. Levy A, Stedman A, Deutsch E, Donnadieu F, Virgin HW, Sansonetti PJ, et al. Innate immune receptor NOD2 mediates LGR5(+) intestinal stem cell protection against ROS cytotoxicity via mitophagy stimulation. Proc Natl Acad Sci U S A. 2020;117(4):1994–2003.

33. Zanello G, Goethel A, Rouquier S, Prescott D, Robertson SJ, Maisonneuve C, et al. The Cytosolic Microbial Receptor Nod2 Regulates Small Intestinal Crypt Damage and Epithelial Regeneration following T Cell-Induced Enteropathy. J Immunol. 2016;197(1):345–55.

34. Lacroix S, Mancassola R, Naciri M, Laurent F. Cryptosporidium parvum-specific mucosal immune response in C57BL/6 neonatal and gamma interferon-deficient mice: role of tumor necrosis factor alpha in protection. Infect Immun. 2001;69(3):1635–42.

35. Theodos CM, Sullivan KL, Griffiths JK, Tzipori S. Profiles of healing and nonhealing Cryptosporidium parvum infection in C57BL/6 mice with functional B and T lymphocytes: the extent of gamma interferon modulation determines the outcome of infection. Infect Immun. 1997;65(11):4761–9.

36. Wolk K, Kunz S, Witte E, Friedrich M, Asadullah K, Sabat R. IL-22 increases the innate immunity of tissues. Immunity. 2004;21(2):241–54.

37. Wolk K, Witte E, Wallace E, Docke WD, Kunz S, Asadullah K, et al. IL-22 regulates the expression of genes responsible for antimicrobial defense, cellular differentiation, and mobility in keratinocytes: a potential role in psoriasis. Eur J Immunol. 2006;36(5):1309–23.

38. Liang SC, Tan XY, Luxenberg DP, Karim R, Dunussi-Joannopoulos K, Collins M, et al. Interleukin (IL)-22 and IL-17 are coexpressed by Th17 cells and cooperatively enhance expression of antimicrobial peptides. J Exp Med. 2006;203(10):2271–9.

39. Zheng Y, Valdez PA, Danilenko DM, Hu Y, Sa SM, Gong Q, et al. Interleukin-22 mediates early host defense against attaching and effacing bacterial pathogens. Nat Med. 2008;14(3):282–9.

40. Pickert G, Neufert C, Leppkes M, Zheng Y, Wittkopf N, Warntjen M, et al. STAT3 links IL-22 signaling in intestinal epithelial cells to mucosal wound healing. J Exp Med. 2009;206(7):1465–72.

41. Parks OB, Pociask DA, Hodzic Z, Kolls JK, Good M. Interleukin-22 Signaling in the Regulation of Intestinal Health and Disease. Front Cell Dev Biol. 2015;3:85.

42. Arshad T, Mansur F, Palek R, Manzoor S, Liska V. A Double Edged Sword Role of Interleukin-22 in Wound Healing and Tissue Regeneration. Front Immunol. 2020;11:2148.

43. Lindemans CA, Calafiore M, Mertelsmann AM, O’Connor MH, Dudakov JA, Jenq RR, et al. Interleukin-22 promotes intestinal-stem-cell-mediated epithelial regeneration. Nature. 2015;528(7583):560–4.

44. Zenewicz LA. IL-22: There Is a Gap in Our Knowledge. Immunohorizons. 2018;2(6):198–207.

45. Lean IS, Lacroix-Lamande S, Laurent F, McDonald V. Role of tumor necrosis factor alpha in development of immunity against Cryptosporidium parvum infection. Infect Immun. 2006;74(7):4379–82.

46. Pollok RC, Farthing MJ, Bajaj-Elliott M, Sanderson IR, McDonald V. Interferon gamma induces enterocyte resistance against infection by the intracellular pathogen Cryptosporidium parvum. Gastroenterology. 2001;120(1):99–107.

47. Barakat FM, McDonald V, Foster GR, Tovey MG, Korbel DS. Cryptosporidium parvum infection rapidly induces a protective innate immune response involving type I interferon. J Infect Dis. 2009;200(10):1548–55.

48. Ayrle H, Mevissen M, Kaske M, Nathues H, Gruetzner N, Melzig M, et al. Medicinal plants--prophylactic and therapeutic options for gastrointestinal and respiratory diseases in calves and piglets? A systematic review. BMC Vet Res. 2016;12:89.

49. Yu JC, Khodadadi H, Malik A, Davidson B, Salles E, Bhatia J, et al. Innate Immunity of Neonates and Infants. Front Immunol. 2018;9:1759.

50. Ashhurst AS, Johansen MD, Maxwell JWC, Stockdale S, Ashley CL, Aggarwal A, et al. Mucosal TLR2-activating protein-based vaccination induces potent pulmonary immunity and protection against SARS-CoV-2 in mice. Nat Commun. 2022;13(1):6972.

51. Girkin J, Loo SL, Esneau C, Maltby S, Mercuri F, Chua B, et al. TLR2-mediated innate immune priming boosts lung anti-viral immunity. Eur Respir J. 2021;58(1).

52. Wu J, Huang S, Zhao X, Chen M, Lin Y, Xia Y, et al. Poly(I:C) treatment leads to interferon-dependent clearance of hepatitis B virus in a hydrodynamic injection mouse model. J Virol. 2014;88(18):10421–31.

53. Wynn JL, Scumpia PO, Winfield RD, Delano MJ, Kelly-Scumpia K, Barker T, et al. Defective innate immunity predisposes murine neonates to poor sepsis outcome but is reversed by TLR agonists. Blood. 2008;112(5):1750–8.

54. Kapoor A, Fan YH, Arav-Boger R. Bacterial Muramyl Dipeptide (MDP) Restricts Human Cytomegalovirus Replication via an IFN-beta-Dependent Pathway. Sci Rep. 2016;6:20295.

55. Secher T, Couturier A, Huot L, Bouscayrol H, Grandjean T, Boulard O, et al. A Protective Role of NOD2 on Oxazolone-induced Intestinal Inflammation Through IL-1beta-mediated Signalling Pathway. J Crohns Colitis. 2023;17(1):111–22.

56. Dalmasso G, Nguyen HT, Yan Y, Laroui H, Charania MA, Obertone TS, et al. MicroRNA-92b regulates expression of the oligopeptide transporter PepT1 in intestinal epithelial cells. Am J Physiol Gastrointest Liver Physiol. 2011;300(1):G52–9.

57. Viennois E, Ingersoll SA, Ayyadurai S, Zhao Y, Wang L, Zhang M, et al. Critical role of PepT1 in promoting colitis-associated cancer and therapeutic benefits of the anti-inflammatory PepT1-mediated tripeptide KPV in a murine model. Cell Mol Gastroenterol Hepatol. 2016;2(3):340–57.

58. Miyamoto K, Shiraga T, Morita K, Yamamoto H, Haga H, Taketani Y, et al. Sequence, tissue distribution and developmental changes in rat intestinal oligopeptide transporter. Biochim Biophys Acta. 1996;1305(1-2):34–8.

59. Tanaka H, Miyamoto KI, Morita K, Haga H, Segawa H, Shiraga T, et al. Regulation of the PepT1 peptide transporter in the rat small intestine in response to 5-fluorouracil-induced injury. Gastroenterology. 1998;114(4):714–23.

60. Ogihara H, Suzuki T, Nagamachi Y, Inui K, Takata K. Peptide transporter in the rat small intestine: ultrastructural localization and the effect of starvation and administration of amino acids. Histochem J. 1999;31(3):169–74.

61. Thamotharan M, Bawani SZ, Zhou X, Adibi SA. Functional and molecular expression of intestinal oligopeptide transporter (Pept-1) after a brief fast. Metabolism. 1999;48(6):681–4.

62. Ihara T, Tsujikawa T, Fujiyama Y, Bamba T. Regulation of PepT1 peptide transporter expression in the rat small intestine under malnourished conditions. Digestion. 2000;61(1):59–67.

63. Foster DR, Landowski CP, Zheng X, Amidon GL, Welage LS. Interferon-gamma increases expression of the di/tri-peptide transporter, h-PEPT1, and dipeptide transport in cultured human intestinal monolayers. Pharmacol Res. 2009;59(3):215–20.

64. Barbot L, Windsor E, Rome S, Tricottet V, Reynes M, Topouchian A, et al. Intestinal peptide transporter PepT1 is over-expressed during acute cryptosporidiosis in suckling rats as a result of both malnutrition and experimental parasite infection. Parasitol Res. 2003;89(5):364–70.

65. Genta RM, Chappell CL, White AC, Jr., Kimball KT, Goodgame RW. Duodenal morphology and intensity of infection in AIDS-related intestinal cryptosporidiosis. Gastroenterology. 1993;105(6):1769–75.

66. Zadrozny LM, Stauffer SH, Armstrong MU, Jones SL, Gookin JL. Neutrophils do not mediate the pathophysiological sequelae of Cryptosporidium parvum infection in neonatal piglets. Infect Immun. 2006;74(10):5497–505.

67. Munoz-Caro T, Lendner M, Daugschies A, Hermosilla C, Taubert A. NADPH oxidase, MPO, NE, ERK1/2, p38 MAPK and Ca2+ influx are essential for Cryptosporidium parvum-induced NET formation. Dev Comp Immunol. 2015;52(2):245–54.

68. Takeuchi D, Jones VC, Kobayashi M, Suzuki F. Cooperative role of macrophages and neutrophils in host Antiprotozoan resistance in mice acutely infected with Cryptosporidium parvum. Infect Immun. 2008;76(8):3657–63.

69. Pavlidis P, Tsakmaki A, Pantazi E, Li K, Cozzetto D, Digby-Bell J, et al. Interleukin-22 regulates neutrophil recruitment in ulcerative colitis and is associated with resistance to ustekinumab therapy. Nat Commun. 2022;13(1):5820.

70. Swale C, Bougdour A, Gnahoui-David A, Tottey J, Georgeault S, Laurent F, et al. Metal-captured inhibition of pre-mRNA processing activity by CPSF3 controls Cryptosporidium infection. Sci Transl Med. 2019;11(517).

71. Lacroix-Lamande S, Mancassola R, Naciri M, Laurent F. Role of gamma interferon in chemokine expression in the ileum of mice and in a murine intestinal epithelial cell line after Cryptosporidium parvum infection. Infect Immun. 2002;70(4):2090–9.

72. Schmidt U, Weigert M, Broaddus C, Myers G. Cell Detection with Star-Convex Polygons. 2018;11071:265–73.

73. Weigert M, Schmidt U, Haase R, Sugawara K, Myers G. Star-convex Polyhedra for 3D Object Detection and Segmentation in Microscopy. 2020:3655–62.

74. Mahe MM, Aihara E, Schumacher MA, Zavros Y, Montrose MH, Helmrath MA, et al. Establishment of Gastrointestinal Epithelial Organoids. Curr Protoc Mouse Biol. 2013;3(4):217–40.

75. Lacroix-Lamande S, Bernardi O, Pezier T, Barilleau E, Burlaud-Gaillard J, Gagneux A, et al. Differential Salmonella Typhimurium intracellular replication and host cell responses in caecal and ileal organoids derived from chicken. Vet Res. 2023;54(1):63.

